# The ribonucleoprotein-mediated CRISPR–Cas9 system induces recurrent Aire gene mutations in contrast to the nickase expression vector in murine *in vitro* or *in vivo* models

**DOI:** 10.1101/2023.10.13.562266

**Authors:** Pedro P. Tanaka, Cíntia J. Monteiro, Max J. Duarte, Ernna D. Oliveira, Ana C. Monteleone-Cassiano, Romário S. Mascarenhas, Mayara C. Vieira Machado, Adriana A. Matos, Letícia A. Brito, Alina O. Oliveira, Thiago M. Cunha, Eduardo A. Donadi, Geraldo A. Passos

**Author notes:** Corresponding author at Molecular Immunogenetics Group, Department of Genetics, Ribeirão Preto Medical School, USP, 14049-900, Ribeirão Preto, SP, Brazil.

## Abstract

Although *in vitro* mTEC cultures can help determine the role of the autoimmune regulator (Aire) gene, the mechanisms of Aire mutations were identified using Aire mutant mice. Nevertheless, long-term cultures of mTECs from mice are difficult to establish. To overcome this, we used a CRISPR–Cas9 system to edit Aire in a murine mTEC line *in vitro* and mouse embryos. Two ribonucleoprotein (RNP) complexes were designed to separately target Aire exons 6 and 8. NHEJ-derived indels or HDR-derived mutations were obtained. Recurrent NHEJ-derived mutations were observed among editions, one in exon 6 (del 3554G) and the other in exon 8 (del 5676_5677TG), i.e., the exon 6 mutation was kept in an mTEC clone edited *in vitro* and *in vivo* in a mouse, and the exon 8 mutation was kept in mTECs *in vitro*. None of the mutations obtained with the nickase system were recurrent, indicating the participation of the RNP complex.

## Introduction

The human autoimmune regulator (Aire) gene is located on chromosome 21q22.3 (1). Mutations in Aire are strongly associated with autoimmune polyendocrinopathy-candidiasis-ectodermal dystrophy syndrome (APECED), also known as autoimmune polyglandular syndrome type 1 (APS1) (OMIM entry # 240300). Most of the knowledge about the Aire gene function in controlling aggressive autoimmunity has been obtained from a mouse model because it is not possible to study this issue deeply in humans (reviewed in 2-4).

The *Mus musculus* Aire gene is located on chromosome 10, position 39.72 cM, is 13 kb long, and transcribes a 14-exon mRNA that encodes a 545 amino acid protein with approximately 77% human-mouse identity (5). To understand the effect of Aire mutations on its biological function and, consequently, the control of aggressive autoimmunity, researchers must address the primary structure of Aire, which encodes a typical domain that interacts with chromatin and regulates transcriptional activity (6, 7).

The monomeric human or mouse AIRE protein has a molecular weight of ∼56 kDa with five domains: the CARD domain (caspase recruiter domain), which is important in the oligomerization of the protein; the NLS domain (nuclear localization signal), which mediates the migration of AIRE from the cytoplasm to the nucleus; the SAND domain (SP100, AIRE1, NucP41/P75, and DEAF1), which promotes protein–protein interactions; and two PHD domains, PHD-1 and PHD-2, which function as histone code readers during chromatin decondensation (8-11).

The human Aire was identified through positional cloning in chromosome 21q22.3, whose genomic region was associated with APECED due to the causative mutations of patients and disease manifestation (1, 12). However, the function of this gene has primarily been validated with Aire knockout (KO) mouse/rat models. Distinct Aire KO experimental models have been reported (13): i) a model with two reproduced mutations observed in APECED patients, one harboring a truncation at the Aire exon 6 that encodes the SAND domain (14) and another presenting a 13 bp deletion in the Aire exon 8, which disrupts the PHD1 domain of the AIRE protein (15); ii) a model of a mutation that reproduces the exon 6 G228W human mutation at the protein SAND domain (16); iii) two models generated irrespective of any known human Aire mutation, one harboring a deletion of the Aire exon 2 (17) and the other with a large replacement from exon 5 to exon 12 of the Aire locus, which yielded a truncated nonfunctional AIRE protein (18); and iv) more recently, an Aire KO rat model harboring a 17 bp deletion in exon 3 that reproduced the R139X human mutation (19).

In addition to the abovementioned models, which were generated through different knockout procedures, a new model of Aire KO mouse medullary thymic epithelial cells (mTECs) was generated by our group through the CRISPR–Cas9 system. This cell line, which is maintained *in vitro* in culture, is a compound heterozygous carrier of indel mutations targeting Aire exon 3 (NLS domain of the encoded protein) and affecting both alleles, which feature different mutations; both alleles produce a truncated nonfunctional AIRE protein (20). Aire KO animals or cells are helpful model systems to investigate the biological/immunological role of the Aire gene/protein. Approaches involving loss-of-function (LOF) have contributed significantly to our understanding of gene function, particularly for mammalian genes. The CRISPR system is among the methods affecting the targeted gene more precisely (21).

The CRISPR–Cas9 system has become a paradigm for genome editing *in vitro* in cultured cells or *in vivo* by microinjecting blastocysts (22-25). However, a given CRISPR system is usually used to edit either cultured cells or embryos.

Here, we evaluated the *in vitro* mutational spectrum of the CRISPR–Cas9 system using an mTEC mouse cell line and an *in vivo* mouse blastocyst embryo model. Additionally, considering that the half-life of the RNP complex is shorter inside transfected cells compared to that of the vector system, we also evaluated the mutation spectrum using an *in vitro* CRISPR–Cas9 RNP complex compared with a CRISPR–Cas9 nickase vector.

## Materials and Methods

### Guide RNA and single-strand oligonucleotide design

The guide RNA (gRNA) or single-stranded oligodeoxynucleotide (ssODN) sequences were designed using the Benchling tool (www.benchling.com). The designs were made according to the following genomic coordinates [Aire, GRCm38 (mm10, *Mus musculus*), location chr10 region 78,030,022-78,043,610 (-), transcript Aire-007], and a 20-nucleotide gRNA was selected. The Cas9-type spCas9 enzyme recognizes the NGG PAM sequence. Exon 6, encoding the AIRE SAND domain, and exon 8, encoding the AIRE PHD1 domain, were chosen as targets. Finally, a list containing the gRNA sequences with their respective on-target and off-target scores and of ssODNs used in the RNP-based system was suggested. The sequences of the top gRNAs or ssODNs used were as follows: Aire exon 6 gRNA (CCAAGAAGUGCAUUCAGGUU), Aire exon 8 gRNA (GGUGAGCUCAUCUGUUGUGA), exon 6 ssODN (CAGGGTGAGCTCATTCAGTCTATCTCTCTCCCAGGAAGATCCAAGAAGTGTATTCAGGTAGGATGGGAGTTTTATACACCCAACAAGTTCGAAGACCCCAGTGGCA), and exon 8 ssODN (AAGAACGAGGATGAGTGTGCCGTGTGCCACGACGGAGGTGAGCTCATCTGCTATGACGGCTGTCCGCGGGCCTTCCACCTGGCTTGCCTGTCCCCACCTCTGCAGGAGATCCCCAG).

For vector-based nickase CRISPR–Cas9 cloning, the 20-nucleotide gRNA was converted to a DNA oligonucleotide with complementary reverse strands with a single guanine added to the 5’ end. Furthermore, the 5’-CACC-3’ and 5’-AAAC-3’ sequences were added to the 5’ ends of the respective forward and reverse gRNA oligonucleotide sequences to allow cloning into a BbsI cleavage site generated in the Cas9n plasmid.

For induction of the HDR pathway during CRISPR-mediated gene editing, a template single-stranded DNA repair oligonucleotide (ssODN) was designed. The 150 bp ssODN was also designed using the Benchling tool. Two point mutations of interest were introduced (a shift of guanine for thymine at position 3561 and a shift of adenine for guanine at position 3563) that alters the amino acid glycine (Gly) for tryptophan (Trp) at position 229 of the Aire protein (Gly229 → Trp229), which in humans corresponds to the G228W mutation. For the PHD-1 domain, a one-point mutation was introduced (a shift of a guanine for an adenine at position 5675) that alters the amino acid cysteine (Cys) for tyrosine (Tyr) at position 313 of the Aire protein (Cys 313 → Tyr 313). Program-suggested sequences were inserted into the ssODN to induce silent mutations in the PAM sequence to prevent gene re-editing.

### mTEC 3.10 cell line

The mTEC 3.10 cells were isolated from C57Bl/6 Aire wild-type mouse thymus. They were maintained through conventional *in vitro* culture in RPMI medium with 10% inactivated fetal bovine serum at 37°C with 5% CO_2_. The mTEC 3.10 cells are phenotypically characterized as CD45^-^, Epcam^+^, Ly51^-^ and UEA1^+^ (19, 26).

### Validation of gRNAs

Conventional PCR using genomic DNA from Aire wild-type mTEC 3.10 cells was performed to amplify exon 6 (254 bp amplicon, primers F: 5’ AGGGCACTGTTTAAGGGT ACA 3’ and R: 5’ TATAGTGACCTGGGCTCCCTT 3’) or exon 8 (392 bp amplicon, primers F: 5’ TCTCTCAGGTGGATGCTGTG 3’ and R: 5’ATGTTATGGCTGCCTTTT GG 3’). The PCR products from each targeted exon were purified and mixed with 0.5 units of recombinant spCas9-GFP (Sigma-Aldrich, Cat # CAS9GPRO), 10 μM gRNA in 2 μL of 10x Cas9 reaction buffer, and RNase-free water in a 20 μL total reaction volume. The mixture was incubated for 60 minutes at 37 °C, followed by 98 °C for 20 minutes for Cas9 inactivation. Reaction products were analyzed through microfluidic electrophoresis in an Agilent Model 2100 Bioanalyzer (Agilent, Santa Clara, CA, USA).

### Cas9 knock-in mouse

The B6J.129(Cg)-Gt(ROSA)26Sor mice were purchased from The Jackson Laboratory (Mouse stock # 024857), mated and maintained at the Animal Facility, University of São Paulo, Ribeirão Preto Campus, São Paulo, Brazil. The Ethics Committee for Research with Animals, Ribeirão Preto Medical School, approved the project (Approval Number 003/2017-1). Our laboratories have biosafety certificate from the National Technical Commission on Biosafety (CTNBio), permit # 0040/98.

### RNP mTEC electroporation

One million Aire wild-type mTEC 3.10 cells were suspended in 100 μL of nucleofector solution (Lonza) containing 20 pmol of spCas9-GFP, 100 pmol of either gRNA and 120 pmol of ssODN. Electroporation was carried out in a Model 2b Lonza Nucleofector using the T-30 program. Then, the GFP^+^ cells were separated into single cells through flow cytometry using a FACSAria Fusion (Beckton, Dickson and Company, USA) apparatus. The cells were individually seeded in wells of 96 flat-bottom plates containing RPMI 1640 medium with 100 pmol SCR7 compound (SelleckChem, USA) for 48 hours at 37 °C with 5% CO_2_ in the atmosphere. Then, the medium was replaced with RPMI 1640 with 10% inactivated fetal bovine serum plus antibiotics. Cell growth was observed every other day. Grown clones were enumerated, individually transferred to wells of 24-well plates and then to 25 cm^2^ culture bottles, and cryopreserved in liquid nitrogen until further analysis.

### Vector-based nickase mTEC electroporation

For Aire exon 6 gRNA (DNA) cloning, the vector pSpCas9n(BB)-2A-GFP (PX461) (Addgene, Watertown, MA, USA, Cat # 48140) was used. The pX461 vector contains the cloning scaffold and the Cas9-nuclease sequence necessary for producing CRISPR–Cas9 complexes and the green fluorescent protein (GFP) DNA sequence. For the insertion of sgRNA in this vector, digestion with the BbsI restriction enzyme (New England BioLabs, Ipswich, MA, USA) was performed. The vector has two BbsI cleavage sites just after the U6 promoter sequence. The digestion removes 22 base pairs, resulting in vector sticky ends, which were used to bind to the gRNA (DNA) that had been added to their ends, complementary bases to these sequences. The mTEC 3.10 cell line was electroporated with recombinant pX461 vectors using the Lonza electroporation kit and the Lonza electroporator, program T-30. The GFP^+^ cells were separated by flow cytometry 48 hours after electroporation using a FACSAria Fusion cytometer.

### Blastocyst microinjection

Four-week-old wild-type C57BL/6 females were superovulated with the hormones PMSG (EMD Millipore) and HCG (Sigma-Aldrich) and mated with C57Bl/6-Cas9-knock-in males. Females with a positive plug were sacrificed, the oviducts were removed, and the embryos were collected and incubated in M2 medium containing hyaluronidase (Sigma-Aldrich, Cat # H3506-500MG). At the one-cell stage, the embryos were washed in M2 medium (Sigma-Aldrich, Cat # MR-051) until all the granulosa cells were removed and then incubated in KSOM medium (Cosmo Bio, Cat # CSR-R-B074) for 30 min at 36 °C with 5% CO_2_ in the atmosphere. The pronuclei of the embryos were microinjected with a solution containing each gRNA plus ssODN. The FemtoJet® 4x series, Femtontip II, and Vacutip I microinjectors (Eppendorf) were used for the microinjection. Embryos were then transferred into the oviduct of pseudopregnant Swiss mouse females, which were crossed with vasectomized Swiss mouse males. Four weeks after birth, the genomic DNA of each pup’s tail was extracted, and then, genotyping was performed through Sanger sequencing of the target exon PCR product.

### Mutation analysis

The Aire exons 6 and 8 that were the targeted genomic regions were amplified through conventional PCR using PlatinumTM high fidelity Taq polymerase (Thermo Fisher, Cat # 11304011) and the oligonucleotide primers as mentioned above.

The PCR products of the targeted regions were sequenced through the Sanger sequencing method. The Fasta sequences of each sample (cell clones or tip-tail genomic DNA PCR products) were analyzed using Synthego’s ICE analysis platform (https://ice.synthego.com/#/), which allows the identification of different types of mutations,

## Results

### In vitro validation of the guide RNAs

The gRNAs designed for exons 6 (SAND domain) and 8 (PHD1 domain) enabled the Cas9 enzyme to cleave the target DNA *in vitro*, as predicted. The respective digestion fragments obtained for each exon were as follows: exon 6 fragments with 130 and 124 bp and exon 8 fragments with 103 and 289 bp (Figure 1).

**Figure 1.**
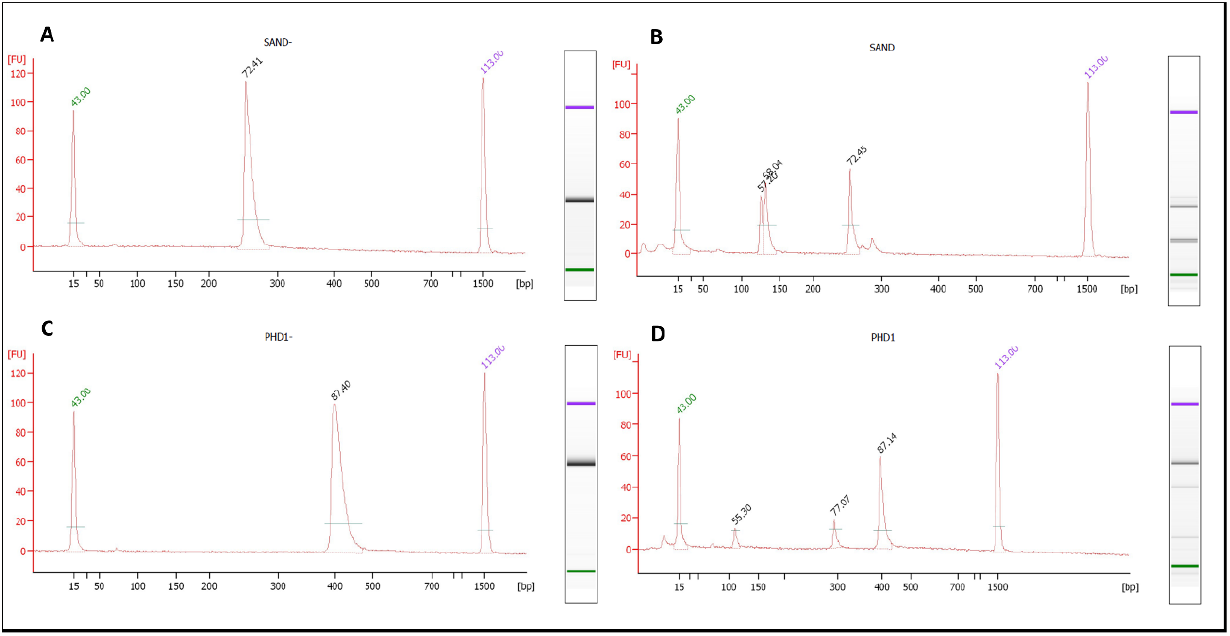
Validation of gRNAs through *in vitro* digestion of target Aire gene exons 6 (SAND domain) or 8 (PDH1 domain) with the Cas9 enzyme. Each exon’s undigested or Cas9-digested PCR products were resolved through microfluidic gel electrophoresis, and the length of the DNA fragments was estimated in base pairs. A) Undigested exon 6, B) Cas-9-digested exon 6; C) Undigested exon 8 and D) Cas-9-digested exon 8.

### mTEC electroporation and single-cell separation

Immediately after mTEC electroporation with RNP-Cas9-GFP or forty-eight hours after electroporation with the nickase-GFP-vector, the GFP^+^ cells were separated into single cells by cytometry using a FACSAria III flow cytometer, and each of the cells was deposited in individual wells of 96-well plates containing RPMI medium with 10% fetal bovine serum and cultured at 37°C with 5% CO_2_ in an incubator atmosphere. The medium was renewed every two other days, and individually grown clones were replicated to 24-well plates and then to 25 cm^2^ culture bottles. Clones were cryopreserved in liquid nitrogen until further analysis.

### Mutant mTEC clones and mice

Figure 2A shows that Sanger sequencing of mTEC clones that were electroporated with the RNP complex evidenced the induction of indel mutations originated by NHEJ and point mutations originated by HDR directed by the ssODN template. The mTEC CS8D6 clone harbors a one base pair deletion that causes a stop codon 51 upstream nucleotide. The mTEC CP5D6 clone has a two base pair deletion that causes a stop codon immediately after the deletions.

**Figure 2.**
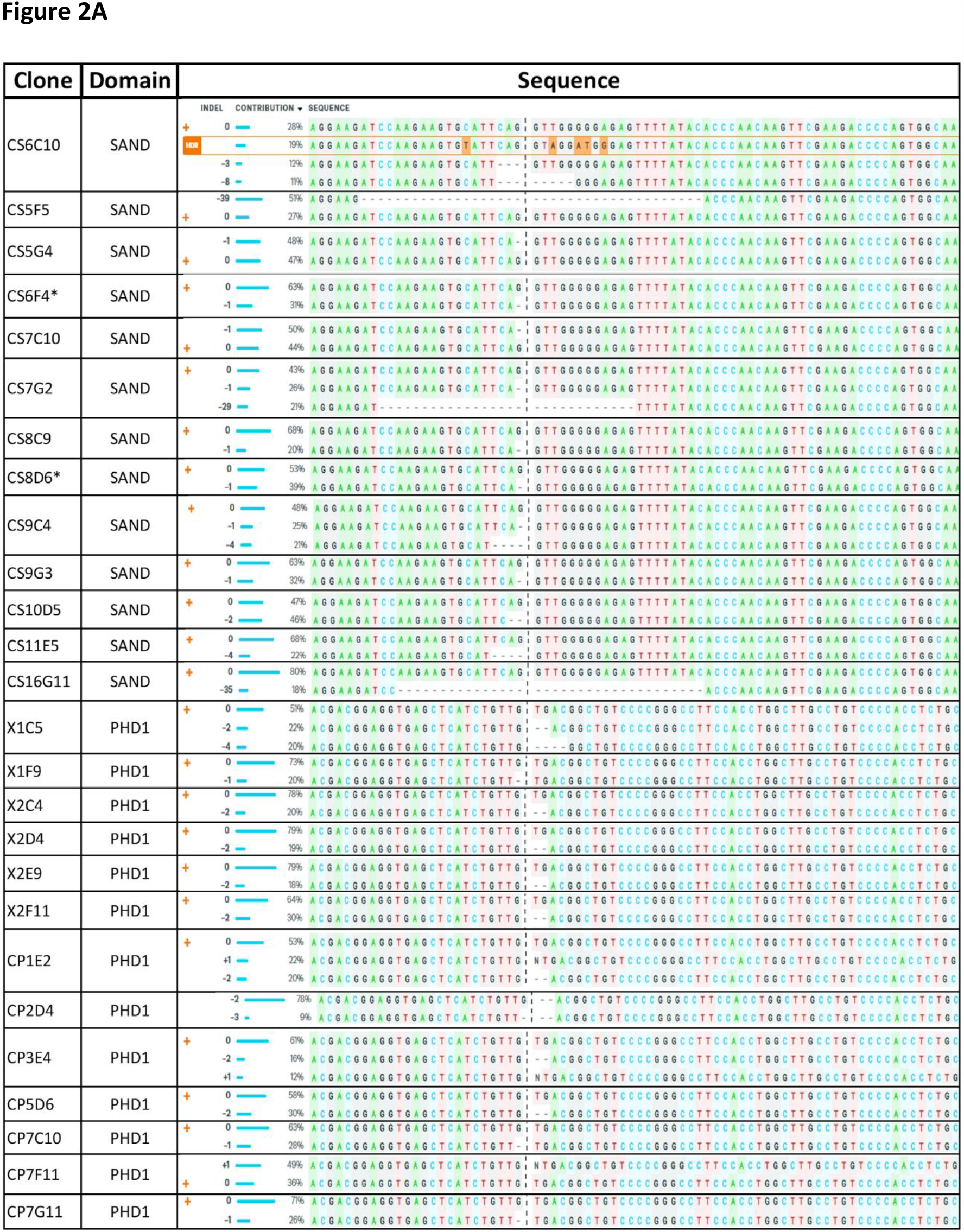
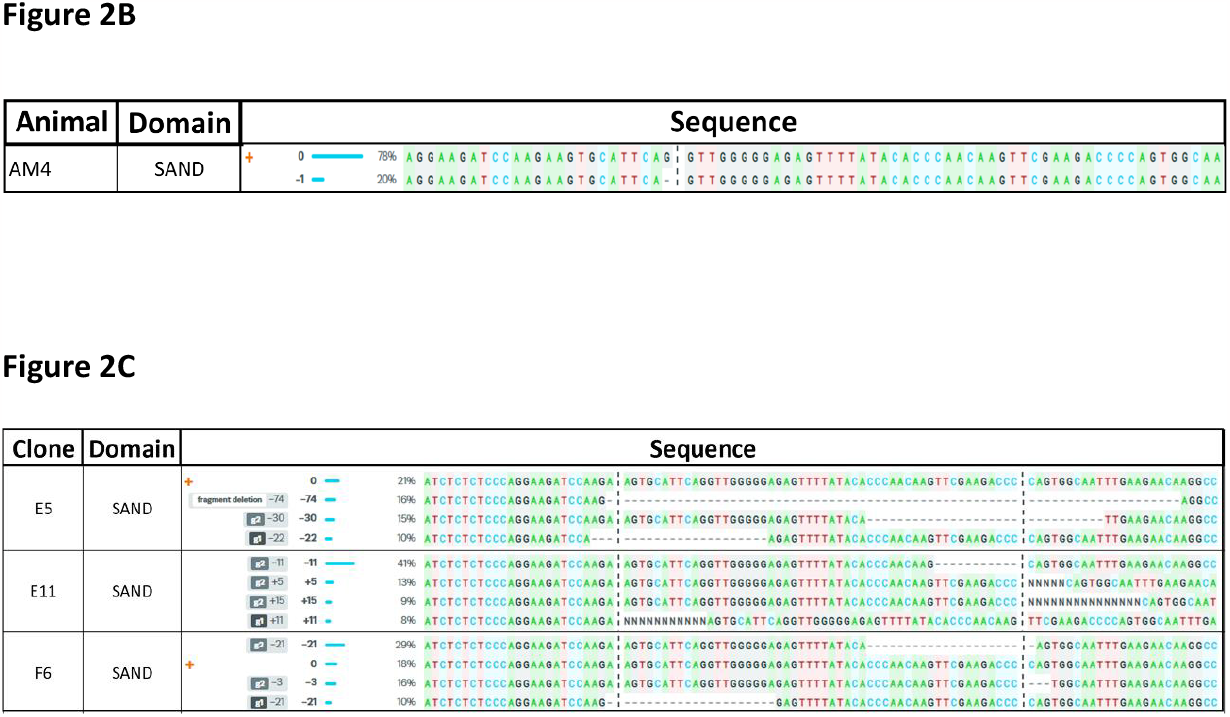
Sanger sequencing of mTEC clones or mice harboring indel mutations. The mTEC cells were electroporated with ribonucleoprotein (RNP) complex or with nickase vector, directed to Aire exon 6 (SAND domain) or exon 8 (PHD1 domain). The mouse blastocyst was microinjected with gRNA directed to Aire exon 6. A) The mTEC clone CS6C10 harbors HDR-induced mutations, clones CS6F4 and CS8D6 (both marked with an asterisk) harbor the same indel mutation as observed in the AM4 mouse (del 3554G), B) Animal AM4 is a heterozygous that harbors the del 3554G in the allele 2, C) mTEC clones E5, E11 and F6 that were obtained by electroporation with the nickase vector exhibited different and nonrecurrent indels in the Aire exon 6.

The exon 6 mutation was recurrent in seven other mTEC clones (CS5G4, CS6F4, CS7C10, CS7G2, CS8C9, CS9C4, and CS9G3), while the exon 8 mutation appeared in eight other clones (X1C5, X2C4, X2D4, X2E9, X2F11, CP1E2, CP2D4 and CP3E4). Among the mTEC clones, the only occurrence observed of HDR-induced mutation, due to the effect of ssODN, was the clone CS6C10 (Figure 2A).

Furthermore, we identified a male mouse whose blastocyst had been microinjected with the RNP complex. This animal showed a recurrence of the mutation observed in mTEC clones CS8D6 and CP5D6 (Figure 2B).

None of the mTEC clones electroporated with the nickase vector showed HDR-type mutations or even recurrence of another type of mutation (Figure 2C).

## Discussion

We investigated the mutational consequences of using an identical CRISPR-Cas9 system of *in vitro* cultured murine medullary thymic epithelial cells (mTECs) and *in vivo* blastocysts to generate edited mice. We targeted the autoimmune regulator (Aire) gene because it is the primary transcriptional regulator in mTECs controlling the expression of a large set of peripheral tissue antigen (PTA) genes, which are essential during the negative selection of autoreactive thymocytes in the thymus that prevents aggressive autoimmunity in the periphery (2, 27, 28).

Mutations in Aire, which in humans maps on chromosome 21q22.3 and in the mouse on chromosome 10C1 (10 39.72 cM) (https://source-search.princeton.edu/), trigger the APECED autoimmune syndrome (OMIM 240300), in which patients display systemic aggressive autoimmunity presenting with adrenal gland failure, dermal dysplasia, and candidiasis, among other complications. Therefore, Aire loss-of-function (LOF) model systems are of great interest in accessing the mechanisms of Aire mutations.

Although cultured mTECs represent an affordable experimental model for Aire (19, 26), the essential aspects of Aire mutations were uncovered using Aire mutant mice (13). Long-term analyses using mTECs harboring a particular Aire mutation, the same mutation found in an APECED patient thymus or an Aire mutant mouse strain, would be interesting. However, isolating and cultivating mTECs from APECED patients or mutant mice is difficult.

A combination of facts allowed the performance of this work, including i) the Aire gene showing significant nucleotide human-mouse sequence identity (∼77%); ii) the possibility of designing gRNAs and ssODNs directed against different murine Aire exons that in humans are involved in APECED; iii) the availability of murine mTEC 3.10 cells, which are cultured *in vitro*; and iv) the availability of B6J.129(Cg)-Gt(ROSA)26Sor, a mouse strain constitutively expressing the Cas9 enzyme.

Then, we used the CRISPR–Cas9 system to mutagenize Aire initially in the murine mTEC 3.10 line *in vitro*. Two guide RNAs (gRNAs) were designed, separately targeting Aire exon 6, encoding SAND, and exon 8, encoding the PDH1 domain of the AIRE protein. In parallel, two single-strand oligonucleotides (ssODNs) were designed to be used with gRNAs as templates for homology-directed repair (HDR) to induce different clones with the mutation G229W in exon 6 and the mutation C313Y in exon 8.

As a result, different mTEC mutant clones were identified, and in most cases, each harbored deletions within the exon 6 or 8 targeted regions. The only HDR-directed mutations identified were in exon 6 (mTEC clone CS6C10).

Of note, recurrent mutations (exon 6 del G3554 and exon 8 del 5676_5677TG) were identified in different mTEC clones and in a mouse harboring the exon 6 del G3554, observed in a mutant male mouse, here termed AM4, which was mated to propagate the mutation.

The CRISPR–Cas9 works with the cell′s DNA repair mechanism to introduce mutations into the target sequence, i.e., after cutting the target DNA by the Cas9 enzyme, the cell activates the template-free nonhomologous end-joining (NHEJ) mechanism, which generates indel-type mutations involving <10 bp (29). Microhomology end-joining (MMEJ) is another repair mechanism that causes larger (>10 bp) indels (30, 31). If the CRISPR–Cas9 system is cotransfected with an ssODN used as a template, the cell can repair its DNA by introducing a mutation in the target (24, 25, 29).

In this study, most of the CRISPR-Cas9-induced mutations that occurred *in vitro* in mTECs originated from the NHEJ mechanism and involved the two Aire targeted exons (exon 6 SAND and exon 8 PHD-1). The use of the SCR7 compound, which is a DNA ligase IV inhibitor, whose enzyme is important for NHEJ (32, 33), seems to have had no significant effect on the model system adopted here, except in one mTEC clone in which an HDR-mediated exon 6 mutation was identified.

Evidence has shown that nonrandom DNA repair may be related to the gRNA sequence/cut region (34, 35) that can explain the mechanism of Aire exon 6 and exon 8 mutations, which in this study were recurrent. In these works, the authors showed that using the same gRNA causes similar mutations even in different cell types. Accordingly, we decided to use an anti-Aire exon 6 gRNA encoded by a nickase vector to assess the involvement of the anti-Aire RNP in the generation of the recurrent mutations.

## Conclusions

Unlike the RNP, none of the mutations obtained with the nickase system were recurrent. Although not studied here in detail, we showed that the RNP complex might have had a more efficient effect on the induction of recurrent mutations.

We generated an array of mTEC clones harboring Aire gene mutations in exons 6 or 8, the exons involved in APECED, and a mouse strain sharing an exon 6 (del 3554G) mutation that can be useful as a model system to assess the role of Aire either *in vitro* or *in vitro*.

## Author′s contributions

PPT, MJD, RSM, ACMC and MCVM: Defined the Aire′s target regions, designed and validated gRNAs, optimization of cell electroporation, singe-cell separation, Sanger sequencing analysis and mutant clone identification; CJM: Conceived the importance of using and designed the nickase vector, raised the importance of mutation recurrence, transfection of cells, Sanger sequencing analysis and mutant clone identification, interpreted and discussed the definitive results and wrote the manuscript; EDO: Defined the Aire′s target regions, designed and validated gRNAs; AAM: Mice genotyping through Sanger sequencing, mice management and crossings; LAB: Cell electroporation, Sanger sequencing analysis, mice genotyping through Sanger sequencing, mice management and crossings; AOO: Blastocyst microinjections, mice managements; TMC: Designed blastocyst microinjection experiments, designed mice crossings and provided the microinjection facility; EAD:

Provided funding and laboratory infrastructure, interpreted and discussed the definitive results and wrote the manuscript; GAP: Conceived the study, provided funding and laboratory infrastructure, supervised the experiments, interpreted and discussed the definitive results and wrote the manuscript.

## Data availability

The datasets used and/or analyzed during the current study available from the corresponding author on reasonable request.

